# MOCHI enables systematic identification of candidate independent intronic transcriptional signals from short-read RNA-seq data

**DOI:** 10.64898/2026.11.05.764473

**Authors:** Dilek Cansu Gürer-Er, Gökhan Karakülah, Daniel Gautheret, Bünyamin Akgül

## Abstract

Standard short-read RNA-seq analysis pipelines quantify expression over annotated exons and disregard signals mapped to introns, thereby missing independently transcribed intronic RNAs. Existing strategies depend on annotation, biologically crude whole-intron quantification, or subjective, low-throughput manual genome-browser inspection. We present MOCHI (**M**apping **o**f **C**ryptic **H**idden **I**ntronic Transcripts), a framework that couples reference-independent, k-mer-based signal discovery with reference-guided genomic interpretation and a custom multi-step filtering module to systematically curate potential independent intronic transcription units from strand-specific short-read RNA-seq data. On two simulated datasets, MOCHI recovered annotated sense-intronic long non-coding RNAs removed from the reference and distinguished them from intronic signal confounders. Across paired ribosomal RNA-depleted and poly(A)-selected libraries from ten cancer cell lines, MOCHI curated candidates whose host introns are strongly enriched for a transcription-initiation signal. Candidate expression is largely uncoupled from host-gene exons, and their genomic features resemble those of annotated long non-coding RNAs. In ten matched cohorts from The Cancer Genome Atlas, candidates were detected in tumor samples, and a subset was associated with overall survival after adjustment for age and host-gene expression. Consequently, MOCHI extends the transcriptomic landscape that can be mined from existing short-read RNA-seq data and provides a curated resource of candidate intronic transcripts for downstream analyses.

## Introduction

Before the advent of high throughput sequencing, introns were known to give rise to two conserved classes of small RNA: small nucleolar RNAs (snoRNAs), most of which are encoded within introns and excised by exonucleolytic processing after pre-mRNA splicing (Kiss, 2002), and mirtrons, microRNA (miRNA) precursors generated from spliced-out intron lariats (Okamura et al., 2007; Ruby et al., 2007). The biogenesis of both classes is, however, coupled to host-gene splicing, so they arise as byproducts of the host transcript rather than as independent transcription units. Whether intronic sequences also harbor their own, independently-transcribed RNAs is not thoroughly investigated.

Large-scale transcriptomic studies have established that the transcriptional output of the human genome greatly exceeds its protein-coding fraction, with the majority of the genome producing reproducible, often cell-type-specific RNA with little or no coding potential (Djebali et al., 2012; Dunham et al., 2012). A substantial part of this signal originates from intronic space. Re-analyses of non-exonic reads have shown that the intronic regions account for the predominant fraction of non-coding RNA (ncRNA) expression in human cells, and that this signal is strand-specific and dynamically regulated (Carninci et al., 2005; St Laurent et al., 2012). Although independent intronic lncRNAs have long been recognized, most intron-derived sequencing signals are routinely attributed to host-gene activity, such as pre-mRNA processing, intron retention, or splicing intermediates, disregarding the possibility of independent, unannotated transcription (Mattick et al., 2023). The GENCODE v7 catalogue formally introduced a *sense_intronic* lncRNA biotype comprising loci residing entirely within protein-coding introns on the same strand (Derrien et al., 2012). Although this categorization was discontinued in 2019 due to the dependence of lncRNA classification on protein-coding gene annotation, the positional information for such lncRNAs is still available from their gene symbol and gene description attributes. In other organisms, stable intronic sequence RNAs (sisRNAs) have been reported as a distinct class of ncRNAs that persists even when splicing is suppressed (Gardner et al., 2012; Jia Ng et al., 2018; Jin et al., 2020). Cryptic transcription start sites embedded within introns are bound by core RNA polymerase II initiation factors and are held silent by repressive chromatin marks, initiating transcription when those marks are removed (McCauley et al., 2021). Several individually characterized intronic lncRNAs, for instance *SLIT2-IT1*, which is silenced by host-promoter methylation independently of *SLIT2* itself, display regulation and function clearly separable from their host genes (Zhang et al., 2022b). More importantly, whether a given base is regarded as exonic, intronic or intergenic is a context-dependent feature of the current annotation protocol rather than an intrinsic feature of the nucleotide itself. Thus, positional, host-referenced nomenclature has been argued to obscure the potential for independent transcription within introns (Mattick et al., 2023). Annotation itself is version-sensitive: transcripts that are intronic under one gene model may no longer be considered intronic under the next, without any change to the underlying sequence (Derrien et al., 2012; Harrow et al., 2012).

Conventional RNA-seq pipelines count reads only over annotated exon features, excluding intronic reads, unless reads have already been annotated as exonic (Liao et al., 2014). Reference-guided transcript assemblers, such as StringTie, Cufflinks, and Scallop partially address annotation gaps by reconstructing transcripts from splice-junction-spanning reads, which is optimized for multi-exon models (Trapnell et al., 2012; Zhang et al., 2022a; Shinder et al., 2026), leading to under-representation of single-exon intronic units. Current annotation protocols, such as the one currently implemented by GENCODE, further compound this bias by excluding single-exon transcripts, unless supported by additional evidence. However, single-exon entities are not necessarily devoid of processing, as such transcripts may still carry a 5′-cap and a 3’-polyA tail. Conventional assembly and annotation approaches, therefore, do not fully capture cell- and condition-specific independent intronic transcripts. Each strategy proposed to recover such transcripts carries limitations that prevent the systematic discovery of unannotated intronic lncRNAs in the human genome. In *Xenopus,* sisRNAs span the entire intron, making their identification relatively easier by quantifying the read coverage of the complete intron (Gardner et al., 2012). However, in humans, this approach aggregates independent intronic transcription with unannotated cassette exons, exon extensions, alternative ends and intron-retention events into a single value indistinguishable from host-gene splicing variation. Peak-calling algorithms, designed for ChIP-seq, such as MACS, have been used to further assess such transcripts in *Xenopus*, as well as in humans (Jin et al., 2020). However, these algorithms collapse the reads from both strands and expect sharp enrichment profiles, which is challenging to attain in human intronic RNAs with dispersed and low-abundance signals. *Drosophila* sisRNAs were discovered and quantified by manual inspection of alignments in a genome browser (Jia Ng et al., 2018), which is subjective, qualitative, relatively irreproducible across users, and more importantly, not scalable to the size and splicing complexity of the human genome. None of these approaches provides a systematic, strand-specific, annotation-independent and isoform-noise-aware framework for discovering new intronic transcriptional units from standard short-read data.

To address these gaps, we developed MOCHI, a framework that couples reference-independent signal discovery with reference-guided interpretation to systematically detect candidate independent intronic transcription units and applied it to cancer as a model system. We hypothesize that conventional pipelines systematically underrepresent independent intronic transcription units and that a reproducible population of such unannotated candidates exists within cancer transcriptomes. To this end, MOCHI is intended as a conceptual framework for reinterpreting intronic RNA-seq signals, complementing the existing novel-transcript-assembly approaches by highlighting reproducible expression signals originating from intronic regions that are not readily explained by transcriptional noise or unannotated splicing isoforms.

## Materials and Methods

### The MOCHI pipeline

MOCHI is implemented as a modular Nextflow DSL2 workflow (v24.10.5) (DI Tommaso et al., 2017). It accepts paired-end, strand-specific short-read RNA-seq data (Figure 1). It integrates a reference-independent, k-mer-based signal-discovery module with a reference-guided alignment module and a custom multi-step filtering framework.

**Figure 1.**
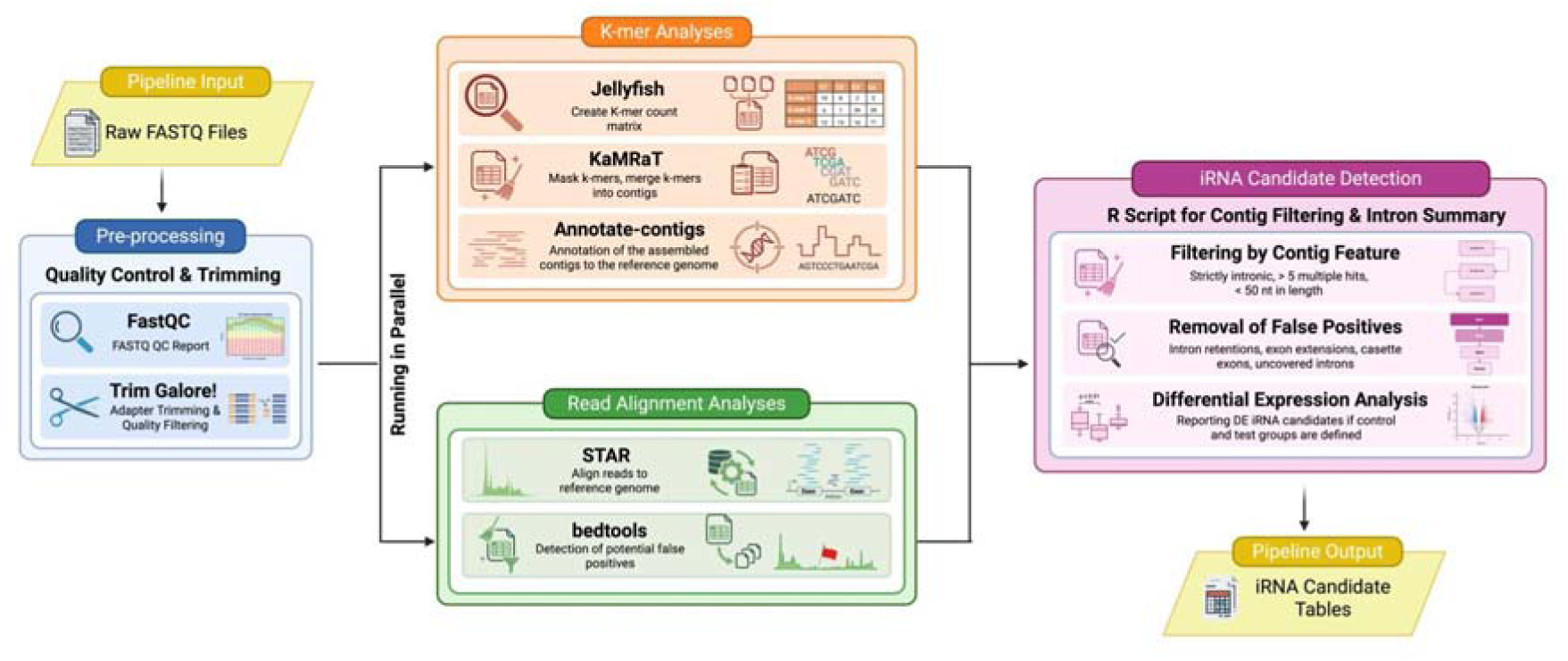
Overview of the MOCHI pipeline. Raw strand-specific FASTQ files are preprocessed (FastQC quality control; Trim Galore! adapter and quality trimming) and then analyzed by two arms running in parallel. The k-mer arm counts 31-mers (Jellyfish), masks transcriptome-matching k-mers, merges the remainder into contigs (KaMRaT), and assigns genomic coordinates and context to contigs (Annotate-contigs). The read-alignment arm maps reads to the reference genome (STAR) and flags potential false positives from intron coverage and splice junctions (bedtools). A custom R module integrates both arms: contigs are filtered by feature (strictly intronic, ≤ few multi-hits, ≥ 50 bp contig length), cleared of false positives (intron retentions, exon extensions, cassette exons, fully covered introns), and, when a control/test design is provided, summarized with differential expression to yield the final candidate iRNA tables.

#### Preprocessing

MOCHI assesses read quality using FastQC v0.12.1 (Andrews et al., 2012) and trims adapter sequences via Trim Galore! (v0.6.10; Phred ≥ 20, minimum length 25 bp) (Martin, 2011; Krueger et al., 2021). To ensure consistent sense-strand k-mer counting, k-mers were reverse-complemented with SeqKit according to library strandedness (Shen et al., 2016) and trimmed reads in original orientation were retained separately for alignment.

#### Reference-independent signal discovery

31-mers were counted from orientation-corrected reads using Jellyfish (v2.2.10) (Marçais and Kingsford, 2011). k = 31 was chosen as the longest odd-length k-mer representable in a 64-bit integer, avoiding palindromic self-complementary k-mers that would confound strand-specific counting. Per-condition count matrices were built with KaMRaT joinCounts (v1.1.1; minimum abundance 5, recurrence set to the maximum replicate number) and indexed (Xue et al., 2024). k-mers matching the GENCODE v47 transcriptome (GRCh38.p14) were removed via KaMRaT mask and the remaining k-mers were assembled into contigs via KaMRaT merge (30–15 overlap). Contigs were assigned genomic coordinates and context (strand; intronic/exonic position; alignment-hit count) with Annotate-contigs (Audoux et al., 2017; El Khaddar et al.) and contigs without valid coordinates were discarded.

#### Reference-guided characterization

Trimmed reads were aligned to GRCh38.p14 with STAR v2.7.11b (Dobin et al., 2013) using GENCODE v47 annotation and permissive splice parameters to maximize sensitivity to non-canonical intronic splice events. Novel splice junctions were retained (≥ 3 unique or ≥ 5 multi-mapping strand-aware reads) and merged across samples. From coordinate-sorted BAMs, bedtools coverage (v2.31.1, split-read, strand-specific) (Quinlan and Hall, 2010) flagged (i) introns covered over > 85% of their length as putative intron-retention events and (ii) > 5 reads within 10-bp exon-boundary windows as putative exon-extension events.

#### Filtering

Candidate refinement was performed in R (v4.4.2) using DESeq2, GenomicRanges, plyranges and biomaRt (Durinck et al., 2009; Lawrence et al., 2013; Love et al., 2014; Lee et al., 2019). Contigs were restricted to intronic/non-exonic positions, and those with > 5 multi-hits or < 50 bp were removed. Remaining contigs were assigned to host introns (canonical transcripts prioritized, with non-canonicals included). Four sequential exclusion filters removed putative retained introns (is_retained), exon extensions (is_extended) and unannotated cassette exons (is_cassette, defined by a novel junction sharing an annotated intron boundary); spliced-within-intron signals (is_spliced) were retained but flagged. Candidates with < 5 reads were discarded. For two-group designs, contig-level differential expression between ribo-depleted (control) and polyA+ (test) libraries was computed with DESeq2 (Benjamini–Hochberg FDR); no fold-change threshold was applied to report all expressed candidates, with their DE statistics. Contig-level results were summarized per host intron (representative = lowest adjusted *P*-value).

#### Implementation and availability

MOCHI accepts a tab-separated design file (sample, condition, replicate), supports pre-built STAR/minimap2 indexes, and can resume from cached intermediates. Dependencies are managed through a conda environment. The pipeline runs on any Linux machine with ≥ 36 GB RAM and a standard multicore CPU; no specialized hardware is required. Runs for this study were performed using the TUBITAK ULAKBIM, High Performance and Grid Computing Center (TRUBA resources).

### Validation on simulated data

Two independent synthetic RNA-seq datasets were generated with polyester (Frazee et al., 2015) to test sensitivity and specificity against intronic confounders. The positive control dataset comprised GENCODE v47 sense-intronic lncRNAs (screened for “PCG-IT#” nomenclature, “sense intronic to” description, and full intronic containment). The negative control dataset comprised synthetically-generated retained introns, exon extensions, alternative first/last exons, and full-length canonical transcripts. The lncRNAs in the positive control dataset were removed from the reference annotation and transcriptome prior to running MOCHI to facilitate their recovery *de novo*. Simulation 1 comprised three confounder categories of 100 regions each and 100 positive control regions (400 regions total); Simulation 2 used a larger, noisier design of 6,165 regions, of which 1,165 were all annotated human sense-intronic lncRNAs. All regions were randomly selected and weighted by the chromosome length to ensure longer chromosomes are represented proportionately. Six samples (3 control, 3 test) were simulated per scenario (strand-specific, error rate 10^-3^, 100-bp reads). Recovered candidates were classified against the positive dataset by strand-aware coordinate overlap using plyranges package (Lee et al., 2019), and precision, recall, accuracy, and F1 scores were computed.

### Dataset curation

Ten cancer cell lines, one per cancer type, were selected to span diverse tissue origins and cellular contexts: HCC827 (lung), HPAF-II (pancreas), MDA-MB-231 (breast), OVCAR-3 (ovarian), Caco-2 (colon), MeWo (skin), U-937 (leukemia), 22Rv1 (prostate), HuH-7 (liver) and LN229 (brain). Each cell line has a corresponding TCGA cohort for downstream clinical analyses. Publicly available human RNA-seq runs were retrieved through the European Nucleotide Archive (ENA) Portal API, filtering for Illumina sequencing, a minimum depth of 30 million reads and a public date from 2012 onward. The following criteria were sought in the cell line datasets: (i) the use of untreated cells; (ii) the use of rRNA-depleted (ribo−) and polyA-selected (polyA+) libraries with at least three biological replicates per library type, and (iii) the use of stranded library preparation protocol, which was confirmed using the RSeqQC package (Wang et al., 2012). Raw FASTQ files were downloaded from the SRA and from ENA. Full run accessions are listed in Supplementary Table S1, and library characteristics are summarized in Table 1.

**Table 1.** RNA-seq datasets and candidate counts across the ten cancer cell lines. Replicate counts are shown as ribo−/polyA+.

| Cancer type | Cell line | Matched TCGA cohort | Replicates<br>(ribo−/polyA+) | Candidate<br>iRNAs |
|---|---|---|---|---|
| Liver | HuH-7 | LIHC | 3 / 3 | 2,681 |
| Leukemia | U-937 | LAML | 3 / 3 | 1,301 |
| Brain | LN229 | GBM | 3 / 6 | 774 |
| Prostate | 22Rv1 | PRAD | 3 / 3 | 381 |
| Ovarian | OVCAR-3 | OV | 3 / 3 | 376 |
| Lung | HCC827 | LUAD | 3 / 3 | 327 |
| Pancreas | HPAF-II | PAAD | 5 / 4 | 258 |
| Breast | MDA-MB-231 | BRCA | 3 / 3 | 252 |
| Colon | Caco-2 | COAD | 4 / 3 | 163 |
| Skin | MeWo | SKCM | 3 / 3 | 135 |

### Downstream characterization of candidates

#### Transcription-initiation support

FANTOM5 CAGE data (robust peaks and total counts, strand-separated) (Forrest et al., 2014; Noguchi et al., 2017) were assessed at the host-intron level in two tiers: (i) host introns containing a robust CAGE-peak summit within their boundaries, and (ii) host introns overlapping a strand-matched CAGE signal of > 10 counts. Enrichment was tested against a length- and GC-matched intronic background by a 1,000-iteration permutation test (empirical *P* with Laplace smoothing). CAGE-signal position relative to each contig (upstream, within, or downstream) was recorded.

#### Expression independence from host genes

Pearson correlations were computed between RPKM-normalized expression of candidates and their adjacent 5′ and 3′ exons, obtained by featureCounts on custom SAF regions, built from MOCHI output contigs. Fully covered intronic regions (removed during filtering) were used as the co-varying control. Within-group differences used the Wilcoxon signed-rank test; between-group differences used two-way ANOVA with Šidák correction. To control for the larger intron pool, a per-cell-line bootstrap (1,000 iterations, size-matched) compared candidate–exon correlation to the null distribution (empirical *P* with Laplace smoothing, combined across lines by Fisher’s method).

#### Sequence conservation

Candidate constraint was assessed with phastCons and phyloP scores from the UCSC 100-way vertebrate alignment (GRCh38). Mean per-candidate scores were compared against the length-and GC-matched intronic background used for CAGE and against annotated sense intronic lncRNAs (with their own matched null) as reference sets. Ribo−-enriched and polyA+-enriched candidate subsets were analyzed separately.

#### Genomic features

Repeat content was quantified by intersecting candidates with RepeatMasker (GRCh38) annotations using bedtools intersect (Quinlan and Hall, 2010; Smit and Hubley, 2025), reporting overlap fraction, full/partial coverage, and repeat class/family composition. Positional bias within host introns was computed from the strand-corrected normalized midpoint (0 = 5′, 1 = 3′ splice site); positional bias of host introns within transcripts was computed as ordinal intron rank / total introns (transcripts with ≥ 3 introns). Distributions are shown as pooled metaprofiles with per-line overlays; the contig-length distribution is reported as an additional descriptor of candidate resolution.

#### Expression and survival in TCGA

Because candidates are unannotated and lack stable identifiers, expression was quantified by coordinate-based coverage: per-patient recount3 BigWig files were summarized over candidate coordinates with megadepth and merged into a candidate × patient matrix per cohort (Wilks et al., 2021a, 2021b). Candidates were blacklisted if overlapping antisense exons determined using strand-flipped GenomicRanges overlap or showing high transcriptome homology computed using BLASTn with > 90% identity and > 80% query coverage (Altschul et al., 1990). Candidates with < 10 counts in ≥ 20% of patients were removed and the remaining counts were VST-normalized using DESeq2 and z-scored (Love et al., 2014). Expression heterogeneity was visualized as hierarchically clustered heatmaps. Association with overall survival was evaluated per candidate by univariate Cox regression (*P* < 0.05), retained through 5-fold cross-validation (significant in ≥ 4 folds), confirmed by age-adjusted multivariate Cox regression ( *P* < 0.05), and finally re-evaluated in Cox models additionally adjusted for host-gene expression to define a host-independent set. Kaplan–Meier curves were generated for representative candidates by median-expression using the survminer package (Kassambara et al., 2016).

## Results

### An integrative framework for intronic transcript discovery

A conceptually straightforward strategy for detecting independent intronic transcription is to computationally inspect RNA-seq coverage across introns as the automated counterpart of manual genome-browser inspection. Such per-base coverage can be computed directly from aligned reads or extracted from pre-computed resources. Pre-computed coverage resources, such as recount3, offer a convenient shortcut, providing readily accessible coverage across large collections of RNA-seq datasets (Wilks et al., 2021b). However, they are not ideal for *de novo* discovery of intronic transcription units for several reasons. As coverage from the two strands is combined into a single track, sense-intronic expression cannot be separated from overlapping antisense transcription. In addition, such resources are limited to a predefined collection of datasets (predominantly human and mouse for recount3), so any new experiment, condition or species falls outside their scope (Wilks et al., 2021b). Moreover, extracting per-base coverage across all introns genome-wide is computationally expensive. We therefore reserved such resources for quantifying already-defined candidate regions. Using stranded coverage information, however, would not be ideal either. Coverage profiles are continuous signals, and isolating a discrete transcriptional unit from such a signal requires change-point detection along the intron (Collado-Torres et al., 2017). Human introns show dispersed, heterogeneous and low-abundance coverage profiles. Thus, segmentation boundaries are difficult to set and often require per-dataset parameter tuning (Gaidatzis et al., 2015). To overcome these problems, MOCHI was designed to utilize k-mer contigs in the discovery step rather than actual read coverage signals. Such k-mer contigs provide localized expression signals, so the discovery step does not depend on dataset-or species-specific parameters. Any strand-specific FASTQ input allows k-mer counting, and building the discovery step on this method gives the framework broad adaptability.

MOCHI was fed raw strand-specific paired-end FASTQ files to run two analytical arms in parallel on the same preprocessed reads (Figure 1). The reference-independent arm counted 31-mers, removed those matching the annotated transcriptome, and assembled the remaining condition-specific k-mers into contigs that were then assigned genomic coordinates, recovering transcriptional signals without being constrained by the existing annotation. The reference-guided arm aligned the same reads to the genome to build a per-sample map of intron coverage and novel splice junctions, which was not used to discover candidates, but to identify and remove the confounders. A custom filtering module in R then intersected the two arms: the contigs were restricted strictly to intronic, non-exonic positions and were cleared of putative intron retentions, exon extensions, and cassette exons before being summarized per-host intron. The design involved three approaches to eliminate the limitations of the existing approaches: (i) annotation independence at the discovery step, (ii) strand-specificity and reproducibility by construction, and (iii) the ability to localize signals to sub-intronic contigs rather than spreading them across the whole intron. The pipeline is capable of running on a single sample or a control/test two-group design. In this study, the ribo− libraries served as the control and the polyA+ libraries as the test for each cell line. Differential expression was computed without a fold change cutoff so that candidates of any polyadenylation status would be retained in the candidate list.

### MOCHI discriminates intronic transcripts from splicing confounders on simulated data

To evaluate whether MOCHI recovers genuine independent intronic transcription while rejecting the unannotated splicing events that confound intron-level quantification, we generated two simulated datasets in which annotated sense-intronic lncRNAs were withheld from the reference and therefore had to be recovered *de novo* against a background of retained introns, exon extensions and alternative first/last exons (Figure 2). In Simulation 1 (300 confounder regions, 100 true targets), MOCHI reported 131 candidates, of which 92 corresponded to the withheld, true sense-intronic lncRNAs, recovering 92 of 100 targets (precision 0.70, recall 0.92, accuracy 0.88, F1 0.79; Figure 2A,C–F). In the larger and noisier Simulation 2 (6,165 regions including all 1,165 annotated human sense-intronic lncRNAs), 913 of the reported candidates were true positives (precision 0.93), recovering 913 of 1,165 targets (recall 0.78, accuracy 0.94, F1 0.85; Figure 2B–F). Performance improved across nearly all metrics as the simulated dataset increased in size and heterogeneity, consistent with the requirements of k-mer-based assembly, which has more signal to work with in a larger, more diverse input. These results indicate that MOCHI can separate independent intronic transcriptional signals from retained introns, alternative and extended exons, and other unannotated splicing isoforms residing in intronic spaces.

**Figure 2.**
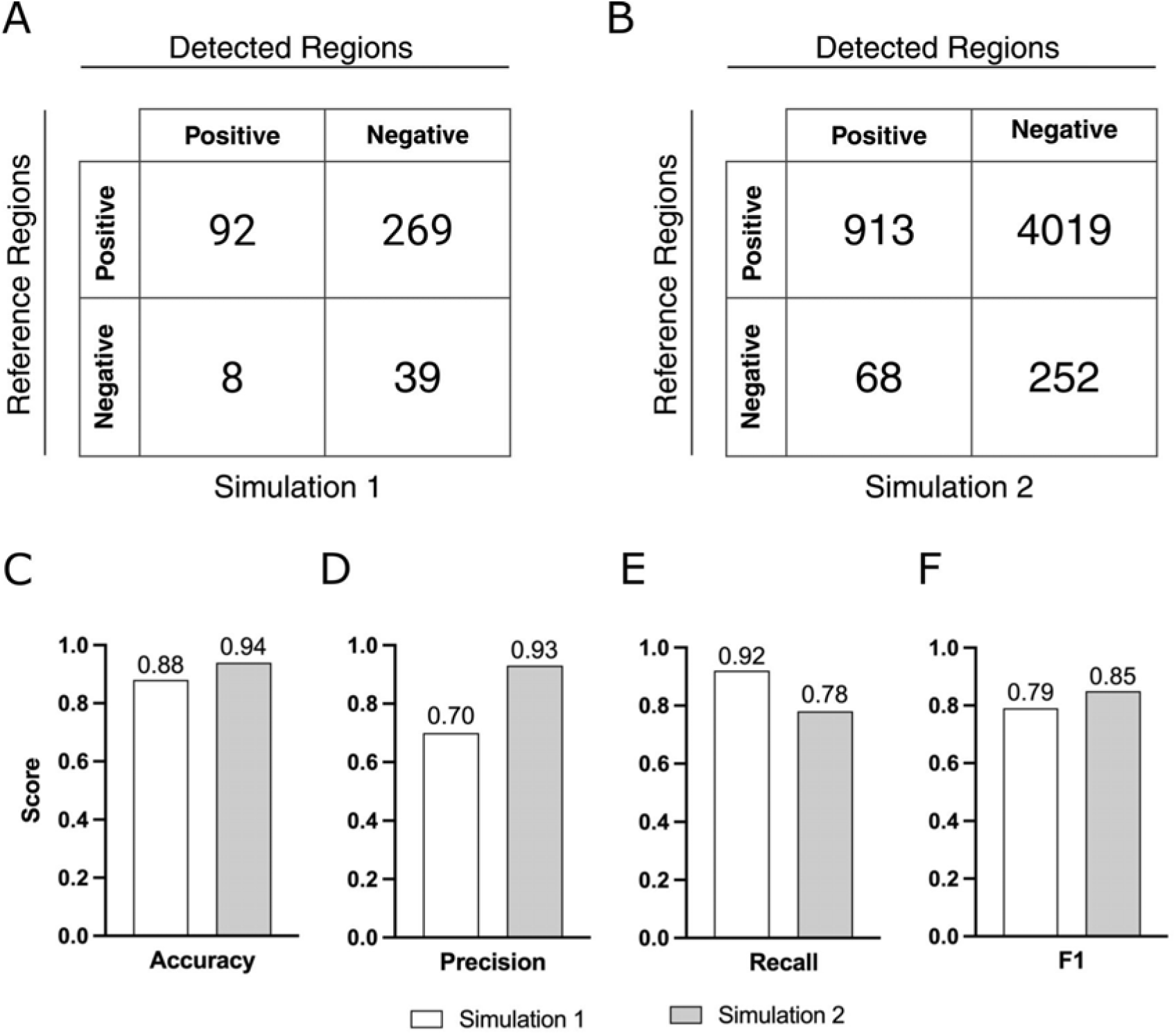
Evaluation of MOCHI on simulated RNA-seq data. (A, B) Confusion matrices of detected versus reference regions for Simulation 1 (300 confounder regions, 100 true sense-intronic lncRNA targets) and Simulation 2 (6,165 regions including all 1,165 annotated human sense-intronic lncRNAs), respectively; positive-class lncRNAs were removed from the reference before running MOCHI. (C–F) Accuracy, precision, recall and F1 for each simulation (Simulation 1: 0.88, 0.70, 0.92, 0.79; Simulation 2: 0.94, 0.93, 0.78, 0.85).

### iRNA candidates are detected across ten cancer cell lines

Applied independently to 20 RNA-seq datasets spanning ten cell lines, MOCHI nominated candidate iRNAs in every case, with per-line counts ranging from 135 (MeWo) to 2,681 (HuH-7) and totaling several thousand candidate loci across the panel (Figure 3A). Coordinates and annotations of all candidate loci, per cell line, are provided in Supplementary Table S2. Sequencing depth spanned ≈ 100–200 million reads for ribo− libraries (median ≈ 164 million) and ≈ 75–200 million for polyA+ libraries (median ≈ 125 million). Ribo− libraries were modestly deeper on average, but no consistent depth advantage was observed across the ten cell lines. With only ten cell lines and 20 independent RNA-seq experiments, a formal depth-count correlation is underpowered, but no obvious trend linked the library depth or the number of replicates to the number of candidates; however, the source of the wide variation in candidate counts cannot be resolved from this analysis.

**Figure 3.**
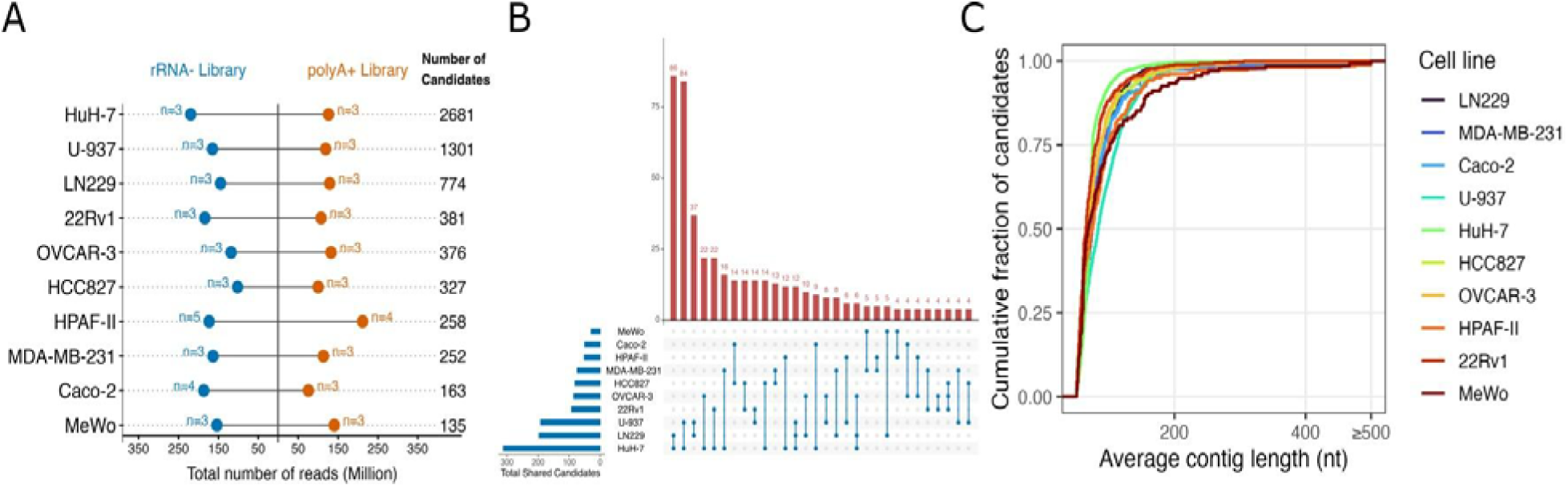
Candidate iRNAs across ten cancer cell lines. (A) For each cell line (labelled by cancer type in the text), total sequencing depth in millions of reads for rRNA-depleted (blue) and polyA-selected (orange) libraries, with biological-replicate counts (n = #), and the total number of candidate iRNAs (right). All expressed candidates are shown regardless of differential-expression status. (B) UpSet plot of candidate sharing across cell lines at the host-intron level (candidate detected in ≥ 2 lines); horizontal bars (lower left) give per-line shared totals and vertical bars the intersection sizes (first 30 intersections). (C) Cumulative distribution of average contig length per candidate for each cell line.

We next asked whether candidates recur across cell lines by mapping them to shared host introns, limiting the comparison by the same host transcript and intron number detected in at least two lines (Figure 3B). Shared loci were concentrated in the three cell lines with the highest number of candidates: U-937 leukemia cell line and HuH-7 hepatocellular carcinoma cell line shared 86 introns, HuH-7 and LN229 glioblastoma cell line shared 84, U-937 and LN229 shared 37, and all three shared 12. Overlaps declined steeply with intersection order, and higher-order overlaps were rare. Intronic transcription detected by MOCHI is therefore predominantly sample-specific, and the candidate set is best regarded as a context-dependent discovery resource rather than a universal catalogue. Because candidates are summarized at the intron level, a single host intron may also harbor more than one distinct transcriptional event. The average per-candidate contig length was short and comparable across lines, reaching a plateau well below the full intron length (Figure 3C). Consistent with the 50-nt minimum-length parameter, most candidates had short average contig lengths, largely in the ∼50-100 nt range (Figure 3C). Each contig therefore marked a localized signal from a region within the underlying transcript rather than the full-length transcript itself, so the contig length reflected the detected signal window and not the transcript’s true extent or boundaries.

### Candidate iRNAs show evidence consistent with independent transcription

We then addressed whether these candidates behaved as independent transcriptional units or as the by-products of host genes. To this end, we sought three orthogonal lines of evidence (Figure 4). First, using FANTOM5 CAGE data as a proxy for transcription initiation, 80.7–88.1% of candidates across cell lines overlapped a strand-matched CAGE signal exceeding 10 counts, whereas only 1.5–5.6% overlapped a FANTOM5 robust peak, and 10.4–17.1% had no detectable CAGE signal (Figure 4A). Because MOCHI does not resolve the exact boundaries of transcripts, CAGE was assessed at the host-intron level. Reassuringly, CAGE signals were located predominantly upstream of the candidate contig, the expected position of a transcription start site relative to a downstream transcript, with within- and downstream-contig signal much rarer (Figure 4B). To account for the limited representation of these cell lines in FANTOM5 and the noisiness of total CAGE counts with respect to robust CAGE peaks, we compared the candidate CAGE-overlap rate to a length- and GC-matched intronic background over 1,000 permutations. Candidate host introns overlapped CAGE signal at ≈ 88% versus ≈ 44% for matched background introns, with no permutation reaching the observed value (empirical *P* < 0.001; Figure 4C).

**Figure 4.**
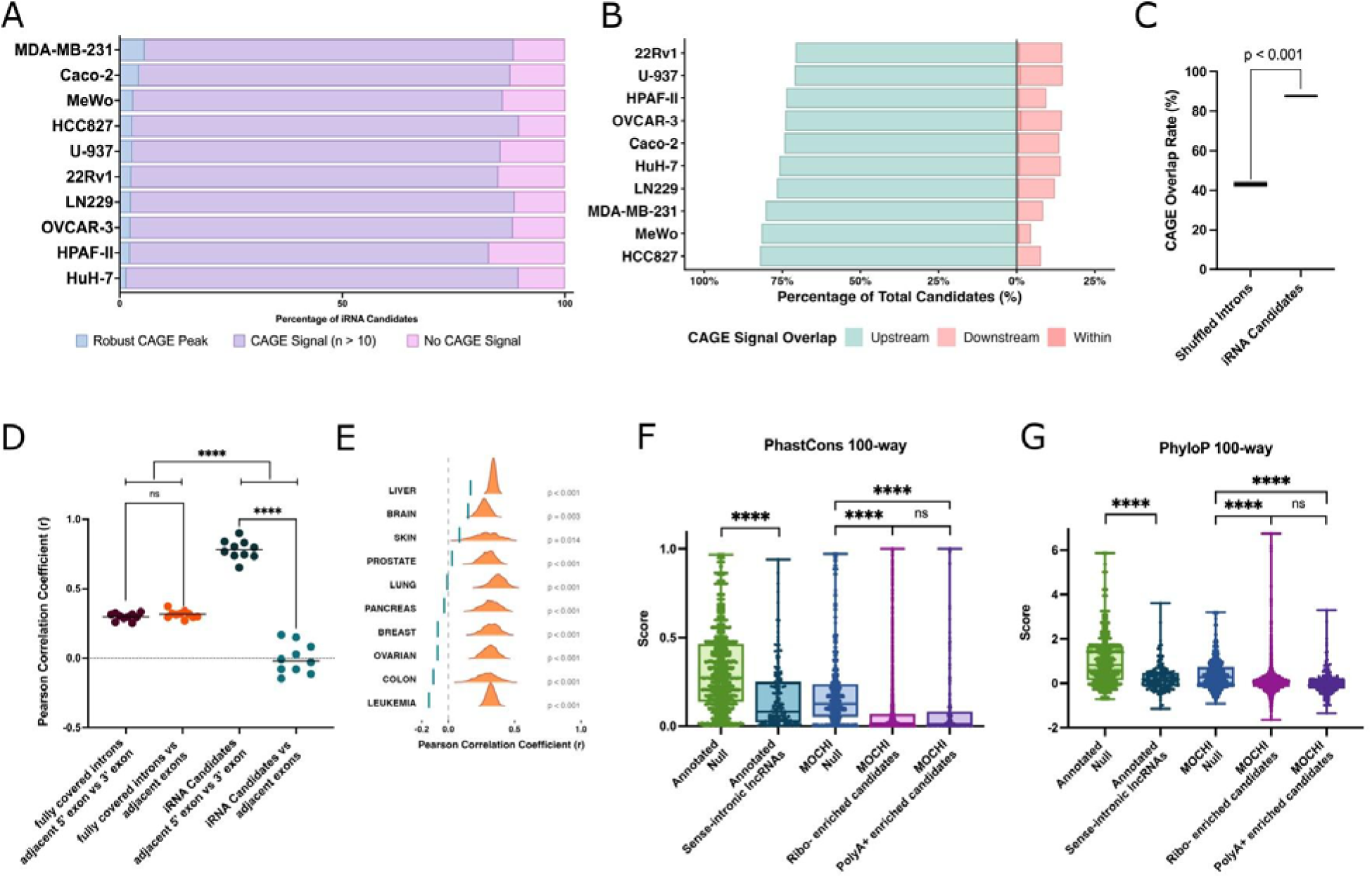
Evidence consistent with independent transcription. (A) Percentage of candidates per line overlapping a robust CAGE peak, a CAGE signal > 10 counts, or no CAGE signal (strand-specific, host-intron level). (B) Position of overlapping CAGE signal relative to the candidate contig (upstream/within/downstream). (C) Permutation test: observed candidate CAGE-overlap rate (≈ 88%) versus length- and GC-matched background introns (≈ 44%; 1,000 iterations; *P* < 0.001). (D) Pearson correlation of expression for fully covered introns and for candidates against their adjacent 5′/3′ exons, and of the flanking exons with each other; each point is one cell line (two-way ANOVA, Šidák correction; **** *P* < 0.0001). (E) Per-cancer-type distributions of candidate–exon correlation. (F, G) phastCons and phyloP (100-way vertebrate) scores for annotated and MOCHI candidate sets against their matched null backgrounds and annotated sense-intronic lncRNAs (**** *P* < 0.0001; ns, not significant).

Second, if candidate reads simply reflected host-gene noise, such as pre-mRNA splicing artifacts or retained introns, their expression would be expected to track the expression of host genes. We therefore compared the correlation between the candidate expression and the adjacent 5′ and 3′ host-gene exons against a control set of fully covered introns removed during filtering. Fully covered introns correlated moderately with their adjacent exons (r ≈ 0.30), similar to the correlation of those exons with each other (r ≈ 0.32; difference not significant), as expected for retention and pre-mRNA signals that co-vary with the host gene. In contrast, although the exons flanking candidates were themselves strongly correlated (*r* ≈ 0.75), candidate expression showed near-zero, slightly positive-to-negative correlation with those same exons (Figure 4D), a difference confirmed by two-way ANOVA with Šidák correction (*P* < 0.0001) and consistent across every cancer cell line examined (Figure 4E). Because the pool of eliminated fully covered introns was substantially larger than the candidate set, we repeated the analysis using size-matched bootstrap sampling in each cell line. Across all ten cell lines, candidate–exon correlations remained below the null distribution generated from fully covered introns and were significant throughout (only 14/1,000 and 3/1,000 bootstrap samples exceeded the observed correlation in MeWo and LN229, respectively; *P* = 0.014 and *P* = 0.003). Candidate expression is thus largely uncoupled from host-gene expression, consistent with the hypothesis of independent transcription.

Third, we assessed evolutionary constraints using phastCons and phyloP scores from the 100-way vertebrate alignment (Figure 4F,G). As a benchmark, annotated sense-intronic lncRNAs, taken as *bona fide* transcripts of this class, were not enriched relative to their matched intronic background. MOCHI candidates likewise showed no enrichment relative to their own background, with no difference between ribo−- and polyA+-enriched subsets. Because a weak constraint is characteristic of genuine lncRNAs and of lineage-specific transcription, the candidates’ low conservation is consistent with lncRNA-like transcription and does not indicate that they are artefacts. We therefore treat conservation as a descriptive feature rather than as evidence for or against a potential function. Taken together, the CAGE enrichment and the pronounced expression uncoupling from host genes provide coherent, if correlative, support that a substantial proportion of candidates represents independent intronic transcription rather than transcription or splicing noise.

### Genomic features of candidate iRNAs

To place candidates in their genomic context, we characterized the repeat content and positional distribution of candidate iRNAs (Figure 5). The fraction of candidates overlapping at least one RepeatMasker element varied widely across the cell lines, the highest in MeWo and the lowest in HuH-7 (Figure 5A), and a notable subset of repeat-overlapping candidates was fully contained within a repeat. At the class level, SINE and LINE elements were the predominant repeats whereas Alu and L1 were the most prevalent at the family level (Figure 5B), mirroring the repeat landscape reported for annotated lncRNAs, in which L1 elements contribute to gene bodies and promoters and Alu elements act as structural modules (Kapusta et al., 2013). More modest, line-specific repeat compositions were also apparent, for example, elevated simple-repeat content in HPAF-II and MIR content in OVCAR-3. Because MOCHI contigs mark only part of a candidate and because transposable-element activation is itself a feature of cancer, fully repeat-covered candidates are reported descriptively. They may reflect repeat-associated transcription rather than *bona fide* independent units, and are retained rather than excluded, consistent with the possibility that some functional lncRNAs are born from transposable-element sequence (Johnson and Guigó, 2014; Burns, 2017).

**Figure 5.**
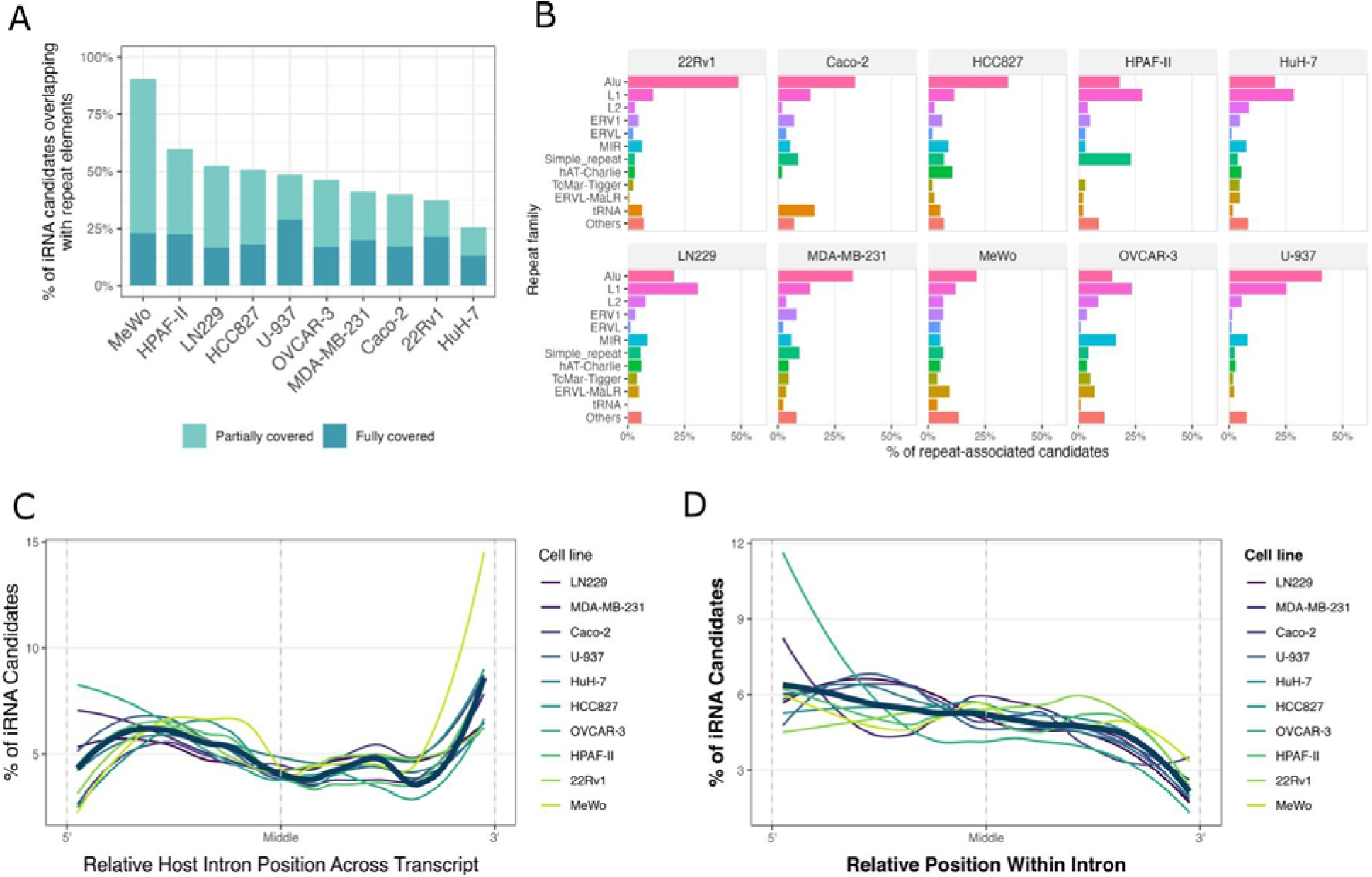
Genomic features of candidate iRNAs. (A) Percentage of candidates overlapping RepeatMasker elements per line, split into partially and fully covered. (B) Repeat-family composition of repeat-overlapping candidates per line. (C) Positional distribution of candidate-containing host introns within parent transcripts (normalized intron rank; transcripts with ≥ 3 introns). (D) Positional distribution of candidates within host introns (0 = 5′ splice site, 1 = 3′ splice site; strand-corrected; per-line smoothed profiles and pooled overlay).

Positionally, candidates were not uniformly distributed within their host introns. Most cell lines exhibited enrichment toward the 5′ splice site with a gradual decline toward the 3′ end, and near-absence of signal at the extreme intron boundaries as expected due to the MOCHI filtering (Figure 5D). At the transcript level, candidate-containing introns were enriched among the 5′-proximal, first introns of their parent transcripts, even after restricting to transcripts with three or more introns (Figure 5C). Because the expression of candidates is uncoupled from that of host-gene exons, this 5′ first-intron bias is compatible with the use of alternative promoters within intronic regions. However, absolute intron-rank patterns must be read together with the relative-rank distribution because they are influenced by differences in transcript architecture across cell lines. Overall, candidate iRNAs are non-randomly distributed, with characteristic repeat composition and positional bias within host introns and transcripts.

### Candidate iRNAs are expressed in primary tumors and associated with survival

Finally, we asked whether iRNA candidates identified in the cell lines are also detected and associated with clinical outcomes in tumors. Because candidates lack stable identifiers, we quantified their expression directly from the genomic coordinates in each corresponding TCGA cohort using per-patient recount3 coverage. Candidates were detectably expressed across cohorts, and hierarchically clustered heatmaps of the top-100 most-variable candidates revealed patient-to-patient heterogeneity within every cohort, showing substantial expression variability (Supplementary Figure S1).

Subsequently, we screened candidates for association with overall survival through a sequential screening approach: univariate Cox regression, 5-fold cross-validation, age-adjusted multivariate Cox regression, and finally additional adjustment for host-gene’s prognostic value (Figure 6A; Table 2). The cross-validation step removed roughly half of the univariately significant candidates in the best-powered cohort (LAML, 73 to 34), underscoring its value in controlling false positives, and the final host-adjustment step further removed candidates whose apparent prognostic value could be attributed to the host locus. After the full cascade, host-independent prognostic candidates remained in all ten cohorts, ranging from 1 (COAD, PRAD) to 22 (LAML) and totaling 61 across the panel. It is important to distinguish between two separate claims. First, the transcript-level independence addresses whether a candidate represents a discrete transcriptional unit rather than host pre-mRNA, intron retention, or transcript extension. This is established through the MOCHI coverage filters together with the CAGE and expression-decoupling analyses described above. Second, the host-expression-adjusted Cox model addresses whether a candidate’s prognostic signal is independent of host-gene expression. The number of prognostic candidates varied markedly across cohorts, with LAML yielding the most. This finding may reflect differences in cohort composition, event rates, and statistical power; LAML is a hematological malignancy and may have distinct coverage characteristics. The observed signal was detected across the cancer-specific cohort analyses but was most prominent in LAML, which we therefore highlight as the primary illustrative example.

**Figure 6.**
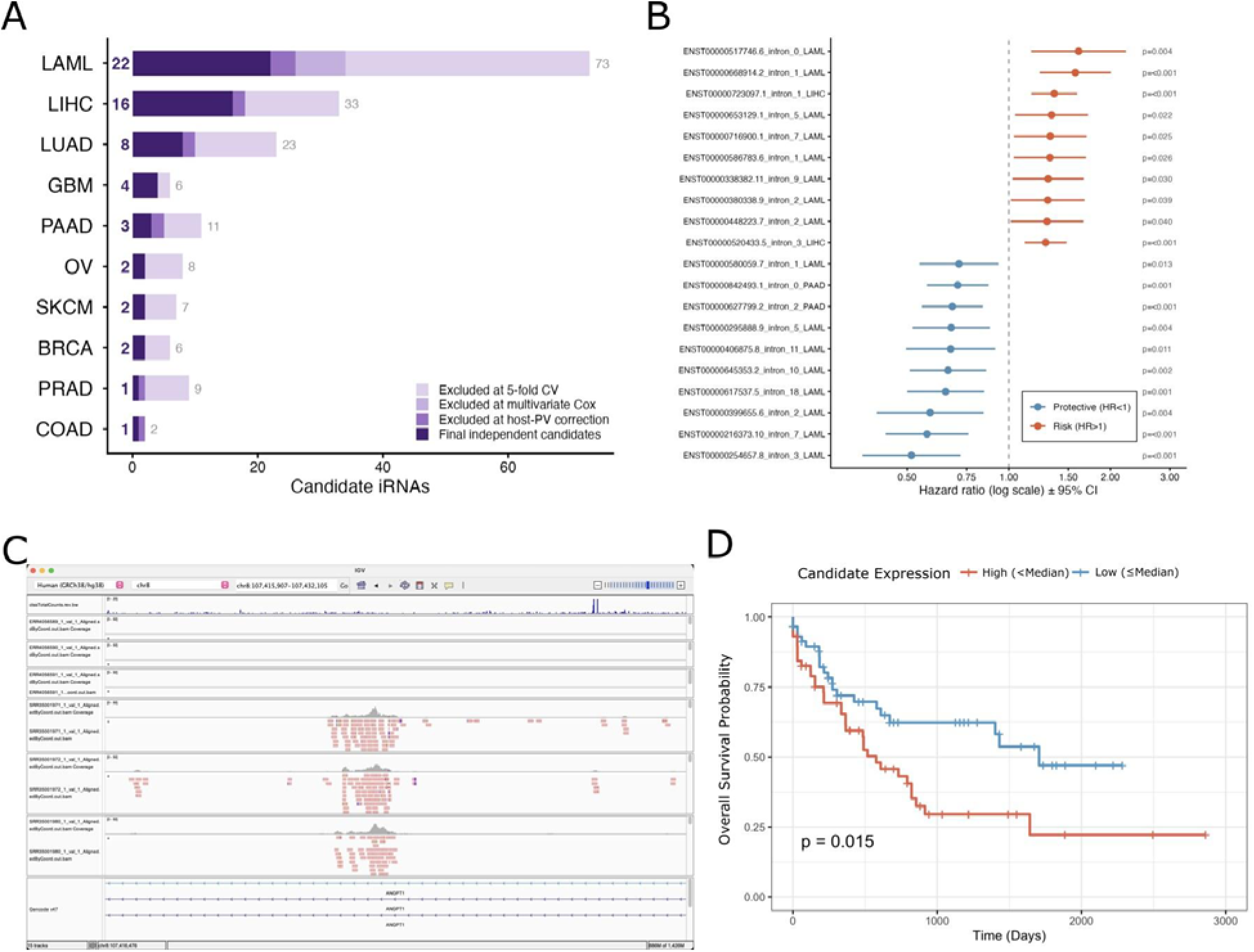
Expression and prognostic association in matched TCGA cohorts. (A) Prognostic-filtering cascade for each of the ten cohorts: candidates are reduced from the univariate-significant set (grey number) through 5-fold cross-validation, age-adjusted multivariate Cox regression and host-gene-expression adjustment to the final host-independent set (bold number; dark segment). (B) Forest plot of representative host-independent candidates (age-adjusted Cox HR ± 95% CI, log scale) coloured by prognostic direction (risk, HR > 1; protective, HR < 1), with *P*-values. (C) IGV view of the *ANGPT1*-intron candidate (chr8) showing localized intronic coverage in polyA+ (SRR) but not ribo− (ERR) samples, with the FANTOM5 CAGE track above. (D) Kaplan–Meier overall-survival curves for the *ANGPT1*-intron candidate in TCGA-LAML, stratified by median candidate expression (log-rank *P* = 0.015).

**Table 2.** Candidate iRNAs retained at each stage of the TCGA prognostic-filtering cascade. Cohorts are ordered by the final host-independent count. Univariate Cox ( P < 0.05) → 5-fold cross-validation (significant in ≥ 4 folds) → age-adjusted multivariate Cox ( P < 0.05) → additional adjustment for host-gene expression (host-independent set).

| *TCGA cohort* | Univariate Cox | 5-fold CV | Multivariate Cox | Final (host-independent) |
|---|---|---|---|---|
| *LAML* | 73 | 34 | 26 | 22 |
| *LIHC* | 33 | 18 | 18 | 16 |
| *LUAD* | 23 | 10 | 10 | 8 |
| *GBM* | 6 | 4 | 4 | 4 |
| *PAAD* | 11 | 5 | 5 | 3 |
| *OV* | 8 | 2 | 2 | 2 |
| *SKCM* | 7 | 2 | 2 | 2 |
| *BRCA* | 6 | 2 | 2 | 2 |
| *PRAD* | 9 | 2 | 2 | 1 |
| *COAD* | 2 | 2 | 2 | 1 |

Across cohorts, the host-independent candidates spanned both risk (HR > 1) and protective (HR < 1) directions (Figure 6B). The most strongly risk-associated candidate across the cancer-specific cohort analyses, in the LAML cohort, lay in the first intron of *ANGPT1* on chromosome 8 (ENST00000517746.6_intron_0; age-adjusted HR = 1.61, 95% CI 1.17–2.22, *P* = 0.004), was detected in polyA+ but not ribo− libraries, carried an upstream CAGE peak, and showed a localized intronic coverage profile without exonic overlap (Figure 6C). Patients with high candidate expression had significantly worse overall survival (median split, log-rank *P* = 0.015; Figure 6D). *ANGPT1* is a known angiogenic factor whose dysregulation has been reported in hematological malignancies (Forster et al., 2011; Xuan et al., 2025). This candidate serves as an illustrative example of the host-independent candidates identified by MOCHI. Collectively, these analyses suggest that MOCHI candidates are expressed in clinical tumor samples and that a curated, host-independent subset potentially carries prognostic information, motivating their further experimental investigation.

## Discussion

MOCHI was designed to recover a class of transcriptional signals that standard short-read RNA-seq pipelines systematically disregard by design. It combines reference-independent k-mer discovery to identify condition-specific signals without annotation bias with reference-guided interpretation to place those signals in their genomic context. This enables independent intronic transcription to be distinguished from the splicing-related confounders that pervade intronic regions. A custom filtering module implements this distinction to identify independent intronic transcriptional units. The result is a hybrid strategy and a curated candidate resource. We emphasize that MOCHI complements existing transcriptome analysis tools rather than replacing them. Transcript assemblers, such as StringTie, Cufflinks, and Scallop are designed to reconstruct full-length, predominantly multi-exon transcripts, whereas lncRNA annotation tools, such as FEELnc, classify transcripts after assembly (Trapnell et al., 2012; Wucher et al., 2017; Zhang et al., 2022a; Shinder et al., 2026). MOCHI addresses a different problem by asking whether localized, unannotated transcriptional activity exists within introns. In doing so, it provides a systematic alternative to manual genome-browser inspection and generates candidate loci for downstream transcript reconstruction and functional characterization.

Several results support the biological coherence of the candidate set while defining the limits of interpretation. The strongest evidence for independent transcription is the combination of CAGE enrichment at candidate host introns (≈88% versus ≈44% in matched background, with predominantly upstream signal) and the near-complete uncoupling of candidate expression from host-gene exons, which stands in clear contrast to the moderate host correlation of the fully covered introns removed during filtering. Both analyses are, however, correlative and constrained by the limitations of short-read data. MOCHI does not define transcript boundaries, so the CAGE support is necessarily assessed at the intron level and should be interpreted as evidence of transcription-initiation activity in the vicinity of candidates rather than a map of the transcript boundaries. The robust-peak overlap is low (1.5–5.6%), and the ≈ 88% figure rests on noisier total CAGE counts from tissues that do not include these cell lines. We therefore keep this piece of information as supportive, not definitive. Conservation analysis showed little purifying selection, comparable to or below the matched background. Given this pattern, we interpret conservation descriptively rather than as evidence for functional relevance. Repeat content is likewise treated as a descriptive feature rather than a filtering criterion. The predominance of Alu and L1 elements mirrors that observed in annotated lncRNAs, yet fully repeat-covered candidates may represent transposable element-associated transcription and are therefore reported separately instead of being excluded.

At the clinical level, coordinate-based quantification showed that candidates are likely to be expressed in tumors and that a host-independent subset associates with survival across cohorts. This analysis carries its own caveats. Candidate iRNA numbers per cohort vary primarily with statistical power and cohort composition. Since the TCGA RNA-seq data were obtained predominantly from the polyA-selected libraries, many ribo−-enriched candidates are likely to be underrepresented in these matrices. Additionally, low death event rates limit power in some cohorts such as COAD. The *ANGPT1*-intronic candidate is presented only as an illustrative example. We deliberately restrict the clinical claim to host-independent prognostic association and do not assert that any candidate drives outcome.

MOCHI users should bear in mind the current tool limitations. First, the absence of transcript boundaries, a primary short-read constraint, motivates targeted long-read RNA-seq of the same lines to resolve candidate termini, as well as 5′/3′ RACE and quantitative PCR for orthogonal validation. Second, the use of one cell line per cancer type limits cancer-specific claims and means recurrence must be read as context-dependent. Finally, functional relevance remains untested. Loss-of-function approaches, together with chromatin-accessibility and histone-modification data, would be needed to extrapolate the nomination to the mechanism of action.

Although MOCHI was tested on cancer datasets, it is not a cancer-specific framework. Cancer was selected as a model system to test the pipeline due to the abundance of publicly available RNA-seq datasets spanning diverse tumor types and experimental conditions, providing a rich resource for systematic method development and evaluation. Cancer is also a biologically favorable setting where non-mutational epigenetic reprogramming, a cancer hallmark, is known to dysregulate chromatin and may increase normally silent intronic transcription to detectable levels, consistent with the pervasive, context-dependent lncRNA expression characteristic of cancer transcriptomes (Iyer et al., 2015; Hanahan, 2022). Consequently, MOCHI provides a general framework for systematic iRNA discovery in any biological context with sufficient transcriptomic data, enabling a more complete characterization of condition-specific transcriptomes by recovering transcriptional signals that conventional RNA-seq analyses routinely overlook.

## Conclusion

MOCHI is an integrative framework that couples reference-independent signal discovery with reference-guided genomic interpretation to systematically nominate candidate independent intronic transcriptional units from conventional short-read RNA-seq data. On simulated data, it recovered known sense-intronic lncRNAs that had been removed from the reference while distinguishing them from intron-retention, exon-extension and alternative-exon confounders, indicating that its candidates are not merely these common intronic artefacts. Applied to ten cancer cell lines, each with paired rRNA-depleted and polyA-selected libraries, MOCHI produced curated candidate sets whose host introns are enriched for CAGE transcription-initiation signal, whose expression is largely uncoupled from host-gene exons, and which show characteristic repeat composition and positional bias; a subset is detectably expressed in corresponding TCGA cohorts and shows exploratory associations with overall survival after adjustment for host-gene expression. Although none of these analyses establishes the candidates as validated transcripts, these candidates provide a prioritized set for targeted experimental validation, and MOCHI itself offers a scalable, reproducible alternative to coarse whole-intron quantification and subjective manual inspection for studying intronic transcription across contexts.

## Data and code availability

All RNA-seq datasets analyzed with MOCHI in this study are publicly available. Full run accessions are provided in Supplementary Table S1. Genomic coordinates and annotations of all candidate regions predicted by MOCHI are provided in Supplementary Table S2. MOCHI is implemented in Nextflow and R and is freely available at https://github.com/dcansugurer/MOCHI under the GPL-3.0 license. A self-contained Apptainer image (v1.0.0), together with the reference files used in this study, is archived at Zenodo (https://doi.org/10.5281/zenodo.23592606).

## Funding

This work was supported by the Scientific and Technological Research Council of Türkiye (TÜBİTAK; Grant No. 122N741 to B.A.) and Ministry for Europe and Foreign Affairs (MEAE) and the Ministry of Higher Education, Research and Space (MESRE) (Campus France; PHC Bosphore; Grant No. 49052ZH to D.G.) within the Bosphorus bilateral cooperation program. D.C.G.E. was supported by the Council of Higher Education (YÖK) 100/2000 Doctoral Scholarship Program, the TÜBİTAK BİDEB 2211-A National Doctoral Scholarship Program, and the French Embassy in Türkiye Research Fellowship Program for PhD students (2023).

## Acknowledgements

The authors would like to thank Şirin Elife Ceren and İpek Erdoğan-Vatansever for valuable discussions and experimental insight that informed candidate prioritization during the development of MOCHI, and H. Atakan Ekiz for constructive feedback on this work. The numerical calculations reported in this paper were partially performed at TUBITAK ULAKBIM, High Performance and Grid Computing Center (TRUBA resources).

## CRediT author statement

**Dilek Cansu Gürer-Er:** Conceptualization, Methodology, Software, Validation, Formal analysis, Investigation, Data curation, Visualization, Writing – Original Draft. **Gökhan Karakülah:** Methodology, Writing – Review & Editing. **Daniel Gautheret:** Conceptualization, Methodology, Resources, Supervision, Funding acquisition, Writing – Review & Editing. **Bünyamin Akgül:** Conceptualization, Methodology, Supervision, Project administration, Funding acquisition,

## Competing interests

The authors have declared no competing interests.

