## Supplemental Figure S1 for "MOCHI enables systematic identification of candidate independent intronic transcriptional signals from short-read RNA-seq data"

### Supplementary Figures

#### Supplementary Figure S1

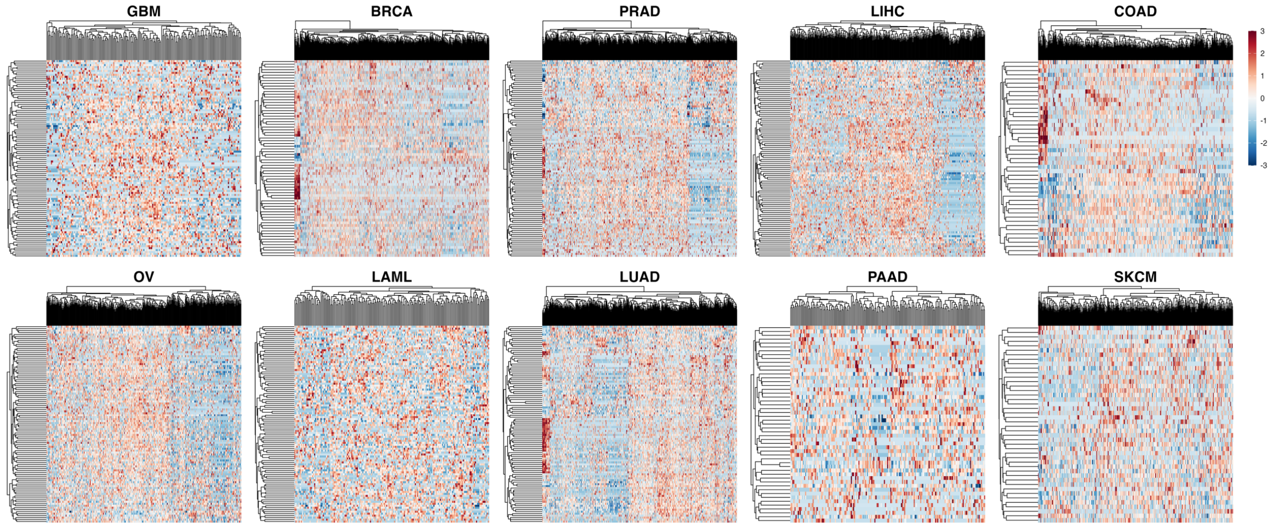


**Expression heterogeneity of candidate iRNAs across matched TCGA cohorts.** Hierarchically clustered heatmaps of the 100 most variable candidates (rows) across patients (columns) in each of the ten matched TCGA cohorts (GBM, BRCA, PRAD, LIHC, COAD, OV, LAML, LUAD, PAAD, SKCM). Candidate expression was quantified from per-patient recount3 coverage over candidate coordinates, filtered ($\geq 10$ counts in $\geq 20\%$ of patients), variance-stabilized with DESeq2 and z-scored per candidate. Color scale indicates z-score (-3 to 3). Rows and columns were clustered by Euclidean distance and complete linkage.
